# Amyloid peptide – synthetic polymer blends with enhanced mechanical and biological properties

**DOI:** 10.1101/2024.07.29.605712

**Authors:** Xianjun Wang, Malay Mondal, Penelope E. Jankoski, Lisa K. Kemp, Tristan D. Clemons, Vijayaraghavan Rangachari, Sarah E. Morgan

## Abstract

Interest in utilizing amyloids to develop biomaterials is increasing due to their potential for biocompatibility, unique assembling morphology, mechanical stability, and biophysical properties. However, challenges include the complexity of peptide chemistry and the practical techniques required for processing amyloids into bulk materials. In this work, two decapeptides with fibrillar and globular morphologies were selected, blended with poly(ethylene oxide), and fabricated into composite mats via electrospinning. Notable enhancements in mechanical properties were observed, attributed to the uniform distribution of the decapeptide assemblies within the PEO matrix. Morphological differences, such as the production of thinner nanofibers, are attributed to the increased conductivity from the zwitterionic nature of the decapeptides. Blend rheology and post-processing analysis revealed how processing might affect the amyloid aggregation and secondary structure of the peptides. Both decapeptides demonstrated good biocompatibility and strong antioxidant activity, indicating their potential for safe and effective use as biomaterials. By evaluating these interdependencies, this research lays the foundation for understanding the structure-property-processing relationships of peptide-polymer blends and highlights the strong potential for developing applications in biotechnology.

## INTRODUCTION

Protein aggregates called amyloids, commonly associated with pathologies such as Alzheimer’s disease, Parkinson’s disease, and spongiform encephalopathies, have been found to form hydrogels at elevated concentrations due to hierarchical self-assembly into large, entangled fiber networks.^1^ Emerging research has uncovered non-pathological roles for amyloids in cellular systems, highlighting their unique mechanical stability and biophysical and biochemical properties.^2–5^ Functional bacterial amyloids, such as the CsgA protein from *Escherichia coli* and the FapC protein from *Pseudomonas*, are examples of amyloids that serve as critical structural components in bacterial biofilms to protect the microorganism.^6–8^ Other than cell-to-cell adhesion^9^ and cell-to-host adhesion,^10^ some functional amyloids and their synthetic analogues have proven to adhere strongly to abiotic surfaces.^11, 12^ Some amyloids also exhibit antimicrobial and/or microbial agglutination activity, where amyloids can entrap pathogens and penetrate their cell membrane, resulting in broad-spectrum antimicrobial properties.^13^ Such remarkable properties of amyloids are associated with their signature cross β-sheet fibril structure in which parallel β-sheets are stacked perpendicular to the fibril axis through hydrophobic interactions, and hydrogen bonds are parallel to the fibril axis.^4, 14^ The strength of a single fibril was reported to be 0.6 ± 0.4 GPa, the same order of magnitude as the strength of steel (0.6 ∼ 1.8 GPa).^4, 15^ Therefore, functional amyloids are attractive building blocks for protein-based functional materials such as drug-releasing vehicles,^16^ tissue engineering,^17–19^ and filtration devices to remove contaminants,^20, 21^ biosensors, ^22, 23^ and bioplastics.^24–26^

A practical aspect of using amyloids as functional materials is developing feasible solution-based processing techniques that retain the desired morphology and properties while removing the solvent. For instance, Li et al. harvested amyloid proteins from plants, partially denatured them, allowed them to self-assemble, and then solution-cast them into thin films. This process led to uncontrolled amyloid nanofibril formation, affecting the transparency of the resulting bioplastic films due to the sensitivity of amyloid morphology to environmental conditions.^24^ Peydayesh et al. fabricated amyloid fibril/polyvinyl alcohol (PVA) blends into free-standing, transparent, and flexible bioplastic films for packaging applications with superior sustainability.^25^ Bagnani et al. investigated mixtures of amyloids extracted from rapeseed cake with methylcellulose and glycerol and reported that the optimized biodegradable plastic blend film was very ductile.^26^ Both works mentioned that the amyloid assembly state (i.e., monomeric or fibril) affects the mechanical properties of composite films. Electrospinning is another popular solution processing technique for developing biomaterials; however, conflicting results were found in fabricating self-assembling peptide/polymer composite nanofiber mats. Rubin et al. found that higher nanofiber peptide incorporation resulted in stronger single fibers. In contrast, Kaur et al. reported that higher peptide loading damaged the bulk mechanical properties of the composite mats.^27, 28^ Due to the complexity of peptide chemistry, it is impossible to directly compare literature reports to summarize the effects of functional amyloids as additives, and we hypothesize that the differences in reported properties are related to differences in the structures and morphologies of the various amyloids evaluated.

Despite advancements in the field of amyloids, understanding the relationship between amyloid sequences, intrinsic self-assembling morphologies, and varied processing strategies for fabricating functional amyloid-based biomaterials remains elusive. The primary challenges include the complex nature of amyloid chemistry and the need for a compatible polymer/solvent system to facilitate amyloid assembly. In our previous publication, we specifically examined a library of *de novo* synthetic decapeptides (DPs) containing an amyloidogenic core.^29^ We systematically varied single amino acid residues in two positions to demonstrate the effects of polarity, hydrophobicity, and charge individually.^29^ Building on this work, we have selected two decapeptides, DP I, and DP II, with well-defined sequences and distinct assembly morphologies for evaluation in the electrospinning of composite nanofiber mats. Pure DP I forms stable β-sheets and assembles into fibril-like structures, which change to a β-turn conformation at elevated temperatures. In contrast, pure DP II exhibits a globular morphology, but its detailed conformation was not fully explored in our previous report.^29^ Polyethylene oxide (PEO) was chosen as the carrier polymer due to its good solubility in the peptide assembly solvent mixture. Different DP/PEO blends were prepared and fabricated into composite mats via electrospinning. The aim of this comparative research effort is to determine the effects of decapeptide sequence, assembly morphology, and loading level on the thermal, mechanical, and morphological properties of composite nanofiber mats. This study lays the foundation for designing new and customizable amyloid-based biomaterials.

## RESULTS AND DISCUSSION

### Secondary Structure and Morphology of Self-Assembled Peptide Aggregates

Two *de novo* peptides, DP I and DP II (Figures 1A and B), were synthesized as described in the experimental section, with crude peptides used without further purification. The purities of DP I and DP II were approximately 75% and 90%, respectively, estimated from LC-MS (Fig. S1 and S2). The two peptide structures were selected due to their propensity to form fibril (DP I) and globular (DP II) morphologies in their purified form, as reported previously.^29^ Peptide structures, FTIR spectra, and AFM morphology images are presented in Figure 1. The deconvoluted FTIR solution spectra show two distinct peaks at 1628 cm ¹ and 1681 cm ¹ for DP I corresponding to β-sheets, while DP II displays a single peak at 1669 cm ¹, indicating β-turns. AFM images of DP I fibrils reveal widths of 20–40 nm and lengths of up to several micrometers similar to those observed previously^29^. DP II forms globular structures with diameters of 15–20 nm (Figures 1E-H).

**Figure 1.**
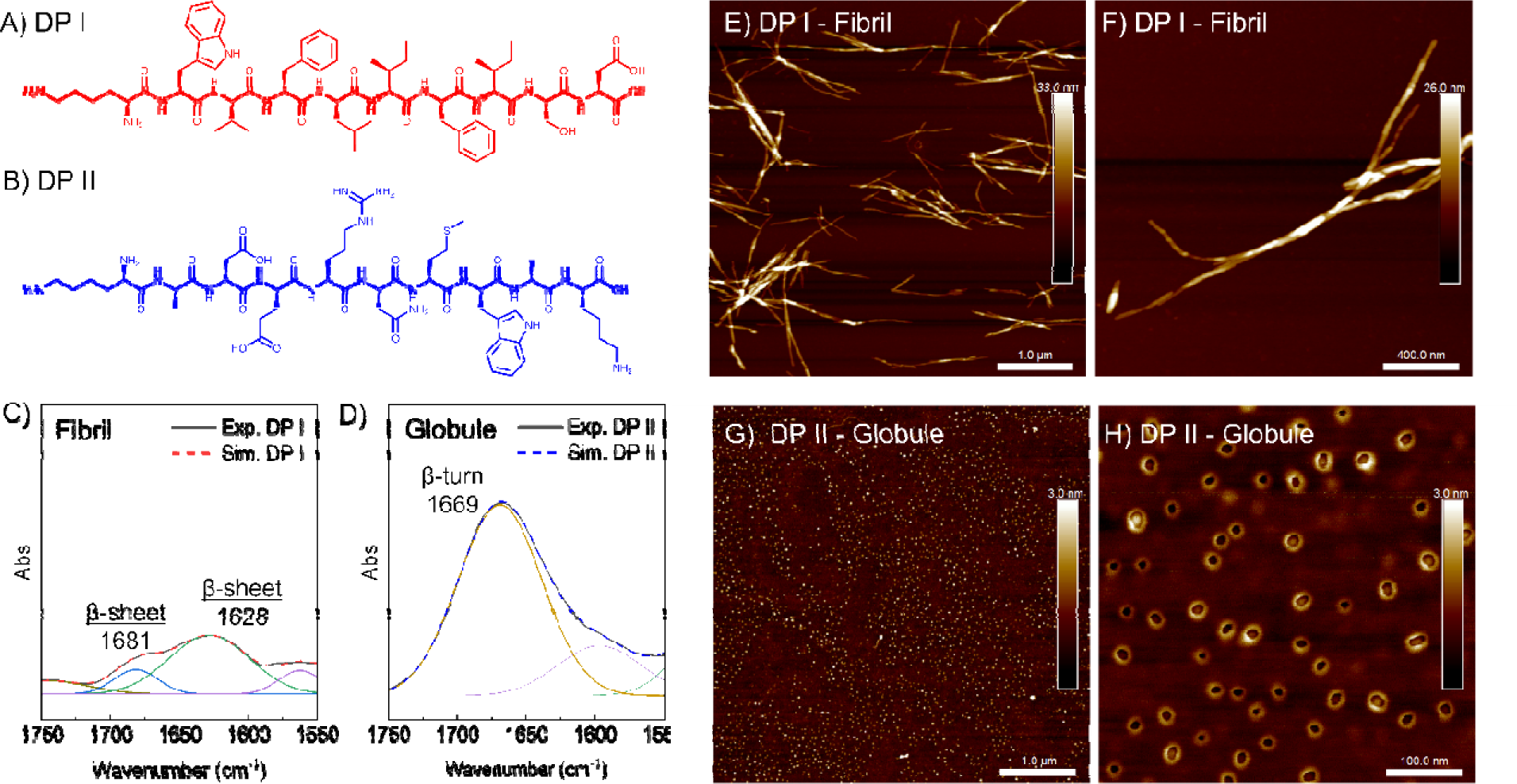
A-B) Structure of DP I and DP II. C-D) Deconvoluted Amide I region of DPs reveals distinct secondary structures, where DP I shows β-sheet character and DP II shows β-turn character. The solid line experimental data and the dashed line simulated data align well. E-G) AFM height images show long fibril-like structures for DPI and globular structures for DPII.

### Morphology of Composite Mats

Solution blends of PEO with varying loadings of DP were prepared and electrospun as described in the experimental section (Table 4). The blends are abbreviated as wt%PEO-DPtype-DP concentration, indicating sample 3-I-1.25 consists of 3 wt% PEO with DP I at a concentration of 1.25 mg/mL. After successful electrospinning, the morphology of the composite mats was characterized using a scanning electron microscope (SEM) (Figure 2), and a detailed fiber diameter analysis is reported in Table 1. Under identical processing conditions, most composite mats display significantly thinner fibers compared to the control (PEO-3), with the exception of sample 3-I-1.25. Increasing DP I incorporation leads to a substantial reduction in fiber diameter, and at higher DP I loading levels (2.50 and 5.00 mg/mL), the composite mats exhibit a much narrower fiber diameter distribution. The loading level of DP II does not affect the mean fiber diameter but narrows the distribution. The observed reduction in fiber diameter and narrowing of the fiber diameter distribution in composite mats may be attributed to shear thinning viscosity effects at electrospinning shear rates (described in the viscosity section) and the zwitterionic nature of DP I and DP II, which alters the conductivity of the polymer/peptide blend. A common strategy to tune the morphology of nanofibers is to add salts to increase the conductivity of the electrospinning solution.^30, 31^ The conductivity of the blank solvent mixture (ethanol/water = 3:1, v/v) is 2.1 ± 0.1 μS/cm. However, the conductivity of the DP solutions increases dramatically, with DP I showing 78.8 ± 0.2 μS/cm and DP II showing 111.2 ± 0.2 μS/cm at a concentration of 1.25 mg/mL (Table SI). Given the intrinsic zwitterionic character of DPs, they enhance the charge density on the surface of the ejected jet during spinning, resulting in higher elongation forces under the electric field. This increase in charge density enhances the overall tension in the fibers due to the self-repulsion of the excess charges on the jet. Consequently, as the charge density increases, the jets become smaller and more spindle-like, leading to a substantial reduction in the fiber diameter.^32^

**Figure 2.**
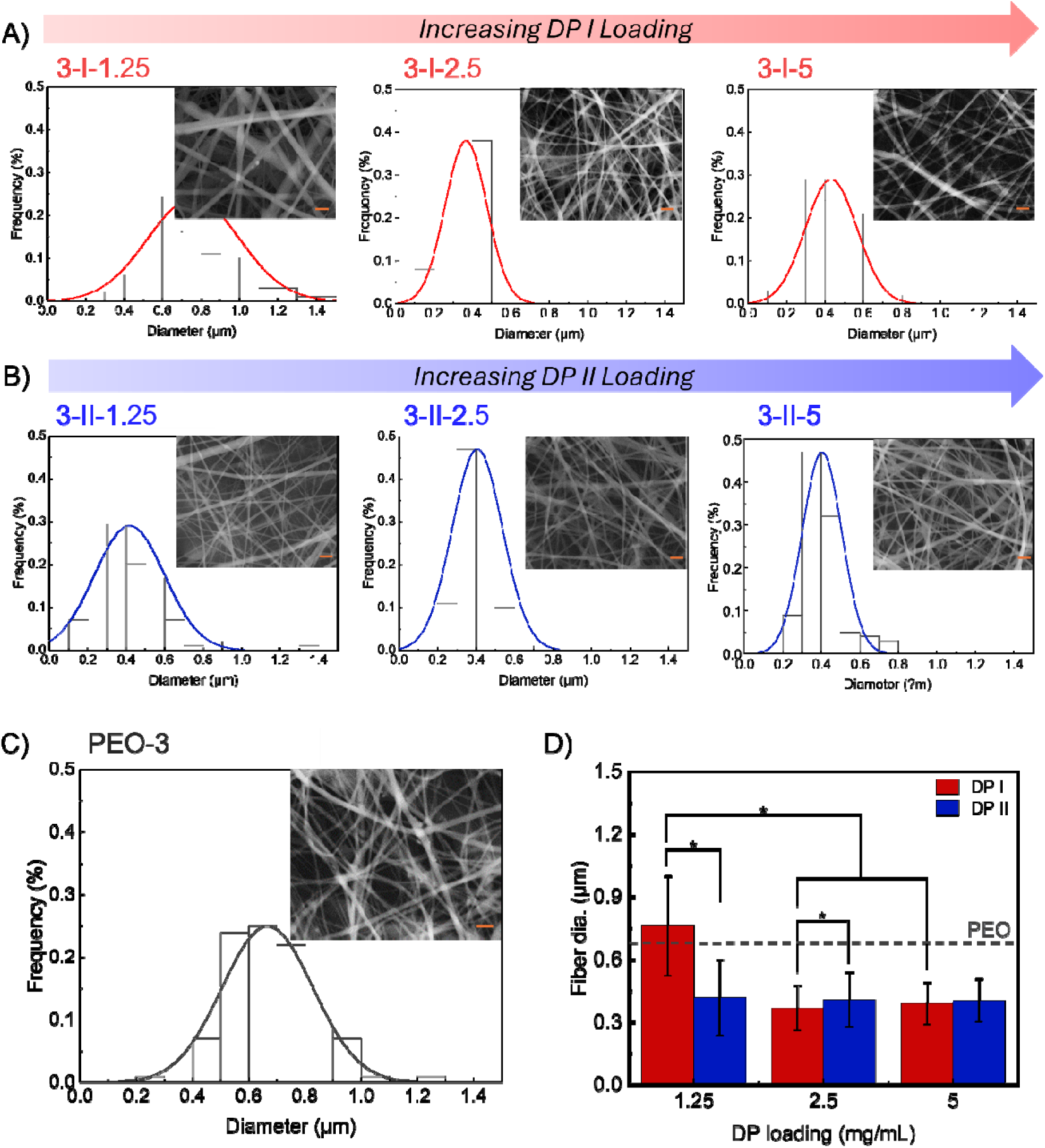
A) Fiber diameter distribution plots of PEO/DP I composite mats. DP I loading level significantly affects the fiber diameter. B) Fiber diameter distribution plots of PEO/DP II composite mats. DP II loading level does not affect the fiber diameter. C) Fiber diameter distribution plots of PEO-3 (the control). D) Fiber diameter plot for electrospun mats. The dashed line represents the control value (PEO-3); * denotes a statistically significant mean difference at *p* < 0.05. All scale bars = 2 μm.

**Table 1.**
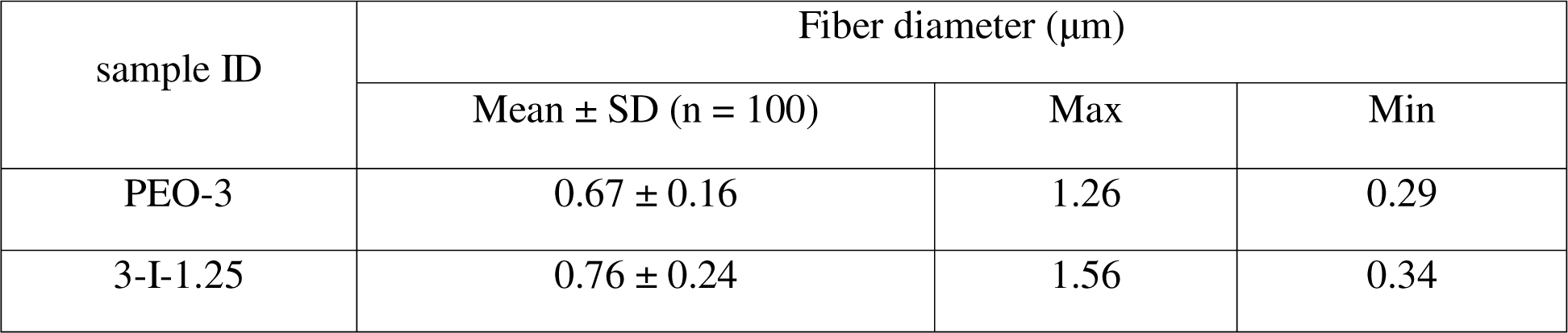
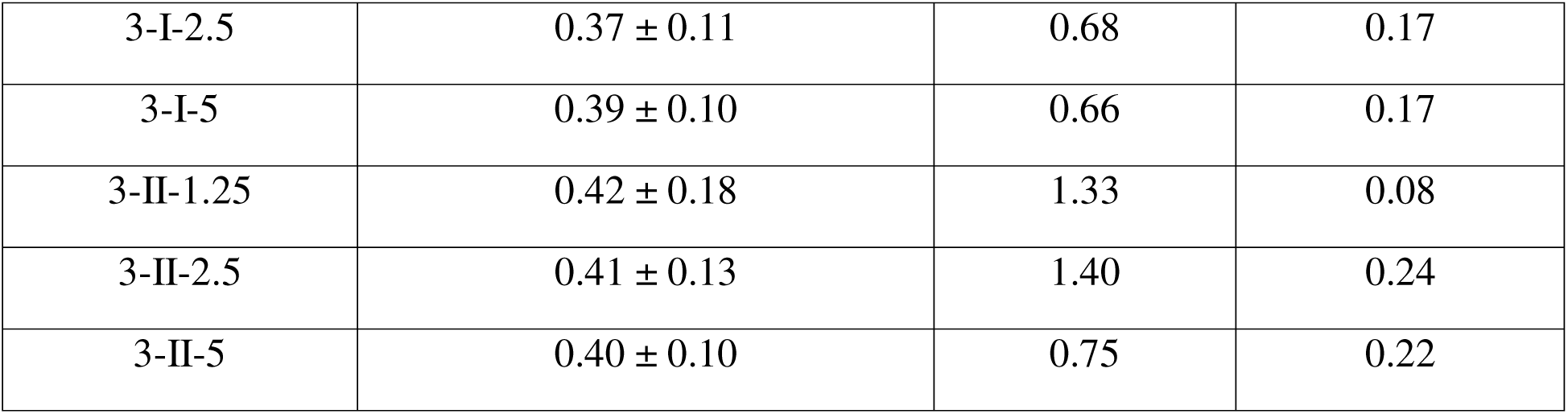
Morphology of PEO/DP composite mats.

### Tensile Properties

We previously reported nanomechanical evaluation of DP I fibrils and measured Youngs’ modulus (***E***) > 3 GPa. ^29^ Using the designed collector (detailed in the experimental section), electrospun mats were fabricated into uniform long strips for tensile testing. The tensile properties of the composite mats are shown in Figure 3 and summarized in Table 2. The tensile properties of the control PEO-3 mat align well with the values reported by Zhou et al.^33^ At all concentrations of DP I and DP II, the average modulus is higher than that of the neat PEO-3 mat (*p* < 0.05), and DP I generally provides a higher modulus than DP II. Notably, the lowest DP I loading level (3-I-1.25) resulted in the highest ***E***, the lowest elongation at break (**ε_b_**), and the largest fiber diameter. With increasing DP I concentration, **ε_b_** increased while ***E*** decreased, but still remained higher than that of neat PEO. This negative correlation between peptide concentration and ***E*** is consistent with findings by Kaur et al, who observed a similar trend with their peptides that formed fibrous networks.^28^ Zhou et al. also reported this behavior for electrospun PEO/cellulose nanocrystal (CNC) composite mats, where increasing CNC loading led to higher ***E*** but lower **ε_b_**.^33^ In our system, we attribute the decrease in modulus as DP I concentration increases to the narrowing of the diameter of the fiber due to increased solution charge. Introducing DP II increased **σ_max_** but did not significantly affect ***E*** or **ε_b_**. A similar pattern has been reported in PEO/ZnO nanoparticle composite fibers by Pittarate et al.^34^ In our system, we attribute the smaller effect of DP II to its globular rather than fibrillar morphology.

**Figure 3.**
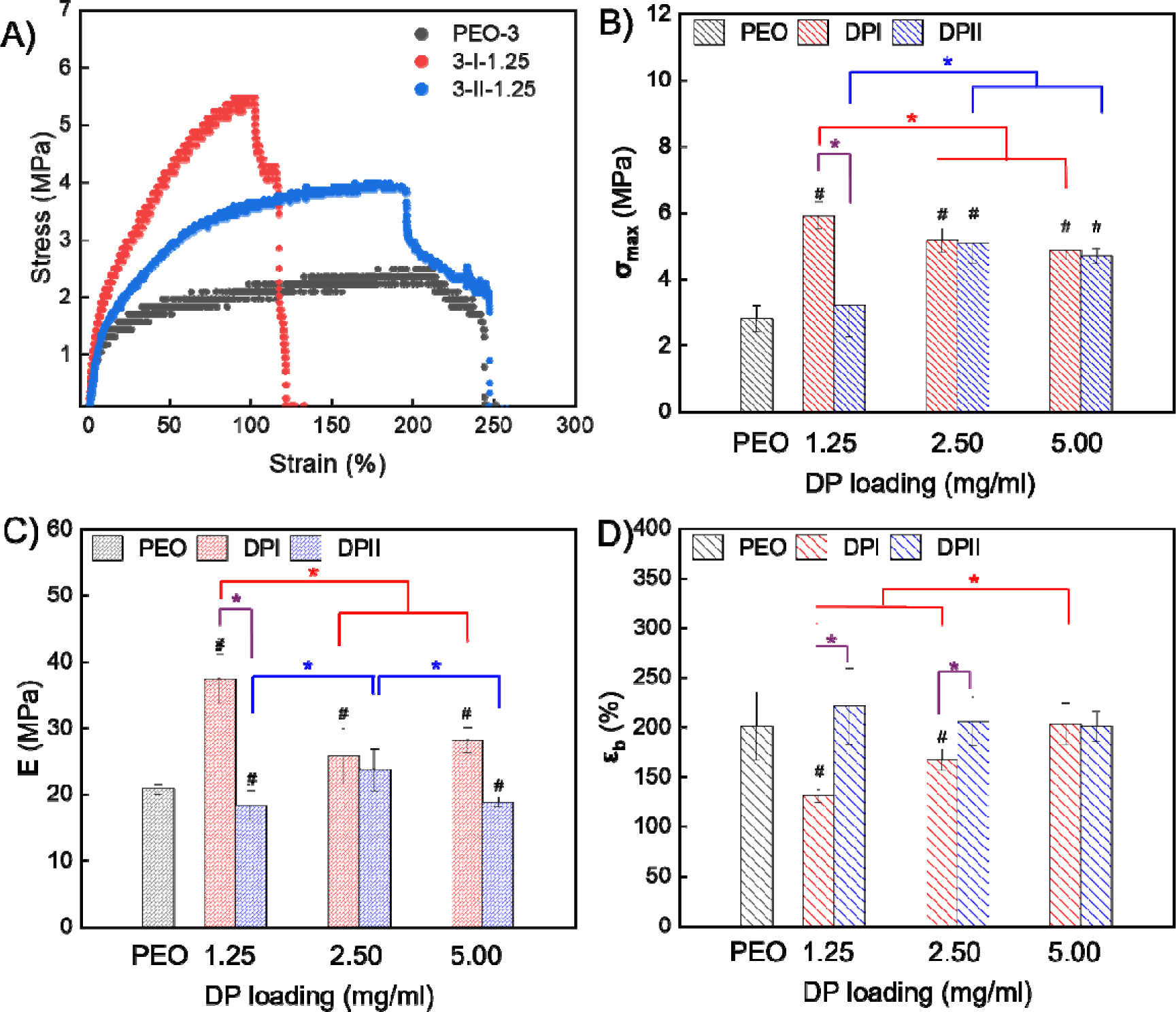
A) Stress-strain curve of samples PEO-3, 3-I-1.25 and 3-II-1.25. B) Comparison of **σ_max_** C) Comparison of ***E*** D) Comparison of **ε_b._** # means statistically different than PEO. Purple bracket: same loading level but different DP; red bracket: DP I at different loading; blue bracket: DP II at different loading; * indicates statistical significance at p < 0.05; Samples collected from two independent runs with triplicates (n = 6). Overall, DP I significantly increases **σ_max_** and ***E*** but reduces ε**_b_** while DP II increases **σ_max_** at high loading but does not affect ***E*** and ***ε*_b_**.

**Table 2.** Tensile properties of composite mats.

| Sample ID | $\sigma_{\max}$/MPa | $E$/MPa | $\epsilon_b$ (%) |
| --- | --- | --- | --- |
| PEO-3 | $2.81 \pm 0.39$ | $20.8 \pm 0.8$ | $201 \pm 34$ |
| PEO (literature) | $2.05 \pm 0.06$ | $15.2 \pm 0.3$ | $200 \pm 12$ |
| 3-I-1.25 | $5.91 \pm 0.42$ | $37.4 \pm 3.7$ | $131 \pm 7$ |
| 3-I-2.5 | $5.17 \pm 0.37$ | $25.7 \pm 4.3$ | $167 \pm 11$ |
| 3-I-5 | $4.86 \pm 0.27$ | $28.2 \pm 1.9$ | $203 \pm 21$ |
| 3-II-1.25 | $3.21 \pm 0.96$ | $18.3 \pm 2.3$ | $221 \pm 39$ |
| 3-II-2.5 | $5.09 \pm 0.62$ | $23.7 \pm 3.2$ | $206 \pm 25$ |
| 3-II-5 | $4.70 \pm 0.24$ | $18.8 \pm 3.2$ | $201 \pm 16$ |

### Blend Viscosity

The apparent viscosity-shear rate relationship of the PEO/DP blends was determined to understand how peptide structure and assembled morphology influence the blend flow characteristics. Figure 4 illustrates that all blends and the control exhibit a yield viscosity at low shear rates, indicating the blend exceeds the entanglement concentration necessary for electrospinning. Adding DP I significantly increases the viscosity of the PEO/DP I blend, whereas DP II does not have a thickening effect. The viscosity-shear rate relationship and yield behavior development are influenced by the aspect ratio and loading level of the two DPs (DP I forms long fibril-like structures with moderate aggregation, while DP II forms well-dispersed nanometer-sized globules).^35, 36^ Higher aspect ratio fillers, like those formed by DP I, percolate at lower loadings due to more efficient network formation, requiring fewer contacts to form a spanning cluster.^37^ DP II may not reach its critical concentration at the levels tested and therefore, the viscosity profiles of PEO/DP II blends are highly overlayed with PEO-3. In electrospinning, the wall shear rate of non-Newtonian fluids is inversely proportional to the inner radius of the needle at a fixed volumetric flow rate.^33^ With an electrospinning flow rate of 0.25 ml/h using a 21G needle, the estimated shear rate is around 50/s. At high shear rates, the viscosity-shear rate relationship becomes independent of blend composition, explaining why the same electrospinning parameters can be used to fabricate nanofiber mats with varying blend compositions. The overall shear-thinning behavior can be attributed to the disentanglement of PEO chains during flow and the disruption of the moderate aggregation of DP I fibril-like bundles.

**Figure 4.**
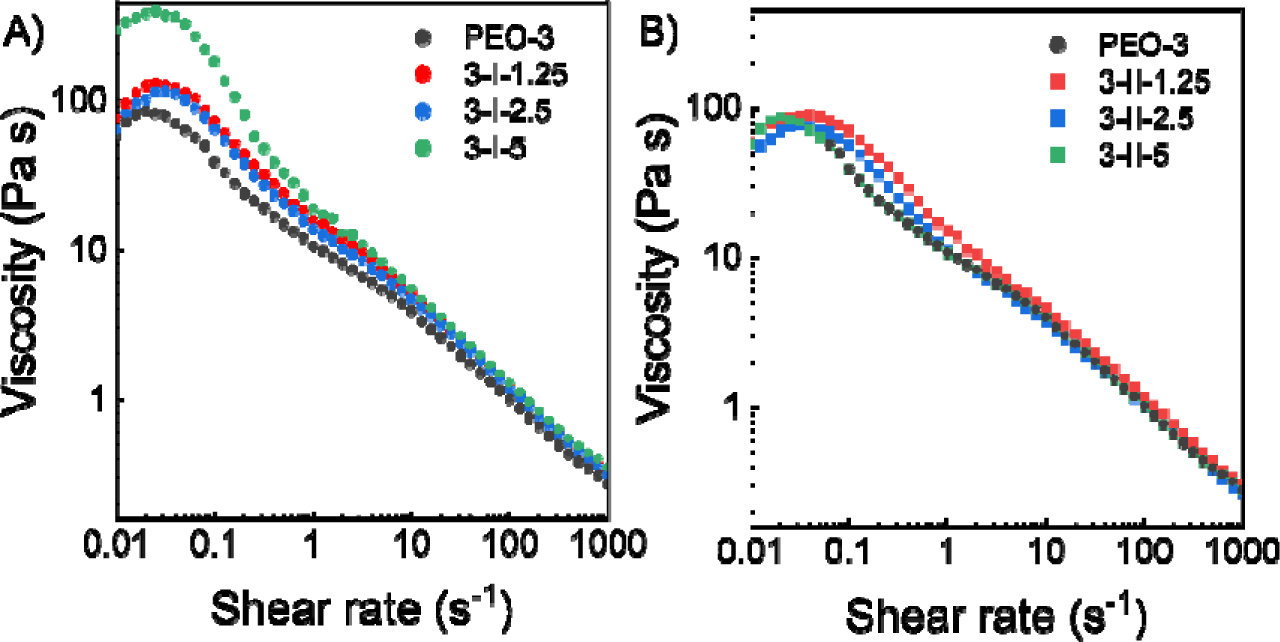
Solution viscosity plots of A) PEO/DP I blends and B) PEO/DP II blends. DP I increases the yield viscosity of the blend and DP II does not have much effect on viscosity.

### Peptide - Polymer Blends and Thermal Properties of Composite Mats

To explain the observation of increased mechanical properties, the peptide incorporation and thermal properties of the composite mats were systematically examined. Figures 5A and B show ATR-FTIR characterization of electrospun mats to investigate whether high voltage and rapid solvent evaporation during electrospinning would disrupt the conformation of DPs. The FTIR spectrum of DP I alone showed a mixture of parallel and antiparallel β-sheet with amide-I bands at 1628 and 1681 cm^−1^(Figure 1C). PEO/DP I composite mats show bands corresponding to parallel β-sheets at1626 and a blue-shifted antiparallel band at 1694 cm^−1^ (+15 cm^−1^) for all blend proportions, with higher DP I ratios showing more intense antiparallel bands (Figure 5A). The observed blue shift is likely due to C=O exciton coupling in an antiparallel arrangement. However, all PEO/DP II mats show a broad peak at 1668 cm^−1^, corresponding to β-turns with no β-sheet signatures. Peaks at 1468 and 1453 cm^−1^ are CH_2_ stretching from PEO.^38^ Together, the data suggest that the peptide secondary structures are preserved after electrospinning with PEO.

**Figure 5.**
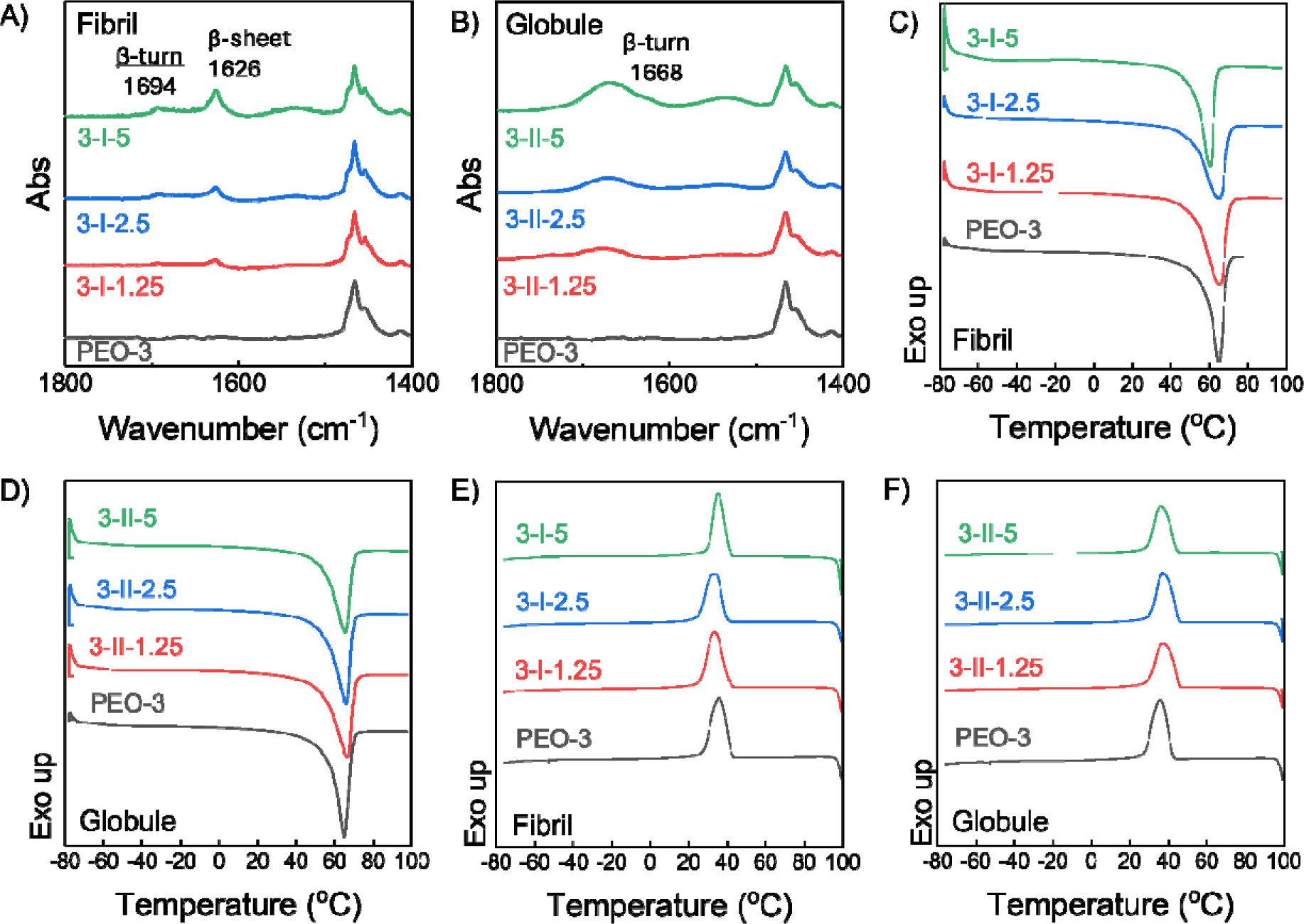
A-B) ATR-FTIR of composite mats showing peptide secondary structures are preserved after electrospinning. C-D) Melting curves from the second heating cycle, and E-F) crystallization curves from the cooling cycle for composite mats compared to PEO-3. Incorporating DP does not dramatically affect the crystallization and melting temperatures but slightly broadens and reduces the melting peak.

The thermal properties of composite mats were characterized using DSC to assess the impact of DP incorporation on the thermal phase transition behavior of PEO. Data is shown in Figure 5 C-F and summarized in Table 3. The electrospun sample PEO-3 exhibited a crystallization peak (*T*_c_) at 36 °C and a melting peak (*T_m_)* at 65 °C with 79% crystallinity, aligning with literature values.^33^ Only small differences in *T_c_* are observed among the composite mats, with values ranging between 33 °C and 37 °C. Specifically, DP I composite mats show a slightly lower *T*_c_ than PEO-3, while DP II composite mats exhibit a slightly higher *T*_c_ than PEO-3. However, these differences are within one to two degrees. Measured *T_m_* is unchanged for the composite mats in comparison to neat PEO, with the exception of the highest loading of DPI, where *T_m_* was reduced by five degrees to 65 °C, suggesting increased PEO chain mobility. Significant decreases in the enthalpy of the melt transition and moderate broadening of the melting peaks are observed for all of the composite mats. The normalized degree of crystallinity is reduced from 79% to ∼60 % for all composite mats, indicating an increase in the PEO amorphous phase. Neither DP I nor DP II exhibit phase transitions between −80 °C and 100 °C. Such behavior is attributed to interactions between the peptides and PEO, and a similar suppressive effect on PEO crystallization was observed in blends of amphiphilic additives (such as fatty acids) with PEO.^39^

**Table 3.**
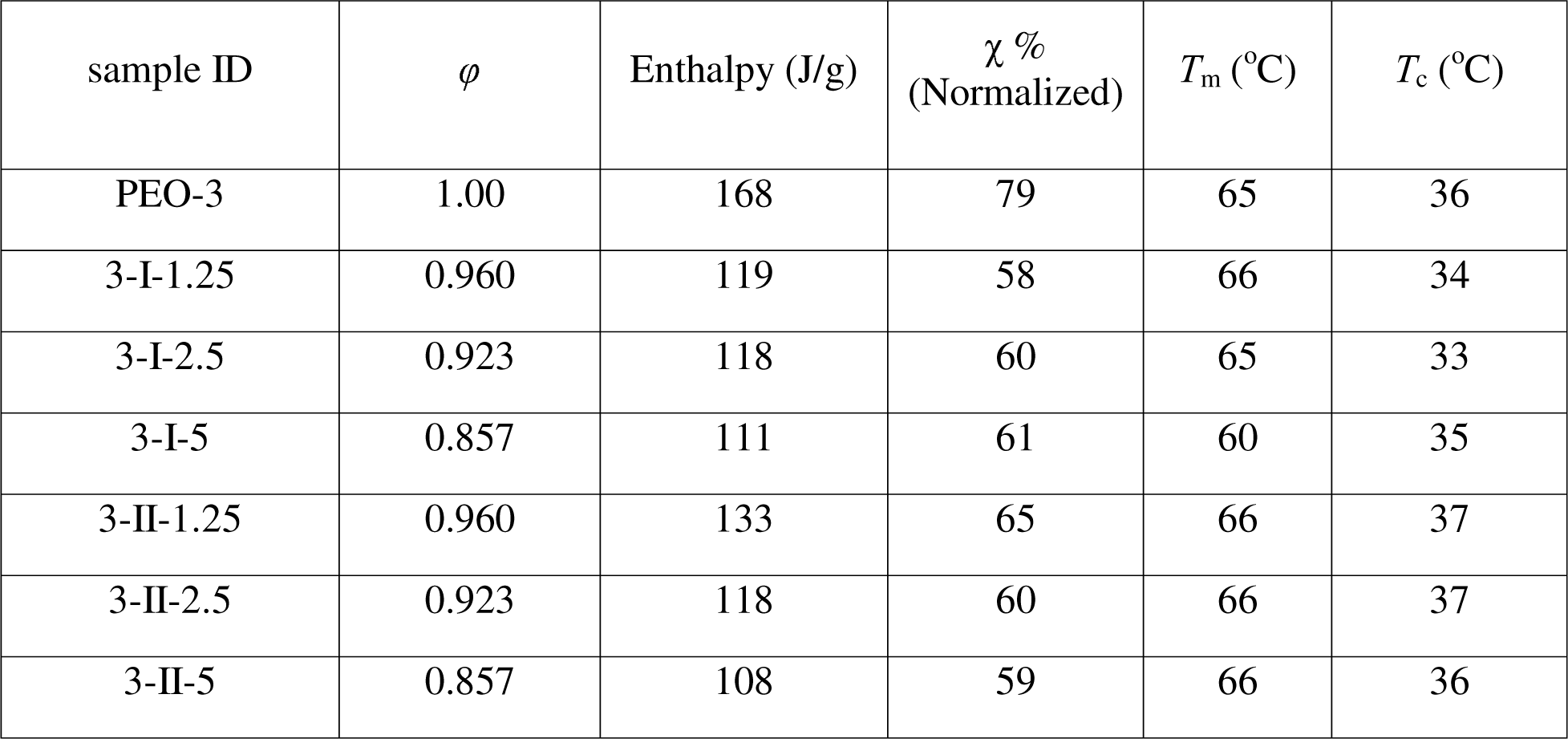
Thermal properties and morphology of PEO/DP composite mat.

### Mesoscale Distribution of Peptides in PEO Matrix

Incorporating DPs in the PEO mats leads to an overall lower degree of crystallinity, and no obvious nanofiber orientation preference was seen in SEM images, which leaves the cause for the mechanical property enhancement in the composite fibers unanswered. To further investigate this system, fluorescently labeled DP I and DP II were co-incubated with unlabeled DPs at two different proportions, blended with PEO and electrospun onto a glass cover slip, and analyzed using fluorescence microscopy. Since the optimal loading level of the labeled peptide after extensive co-incubation was unknown, two levels of fluorescently labeled DPs (0.1% and 0.05%) were screened, and fluorescence images are shown in Figure 6. Overall, both DPs showed good dispersion in the PEO matrix after electrospinning. It is important to note that the glass coverslip is less conductive than the metal collector used previously, so the deposition pattern and fiber morphology observed are expected to differ from those seen in the original SEMs. DP I and DP II have assembled dimensions much smaller than the diameter of nanofibers collected on the glass cover slip. Therefore, under 100x magnification, individual DP I fibrils and DP II globules cannot be distinguished; instead, a blurred fluorescence signal is observed in the PEO matrix. Based on this observation, we attribute the enhanced mechanical properties to the good dispersion of peptides within the PEO matrix, which enables efficient stress transfer, and the fibrillar DPI provides greater reinforcement than the globular DP II.

**Figure 6.**
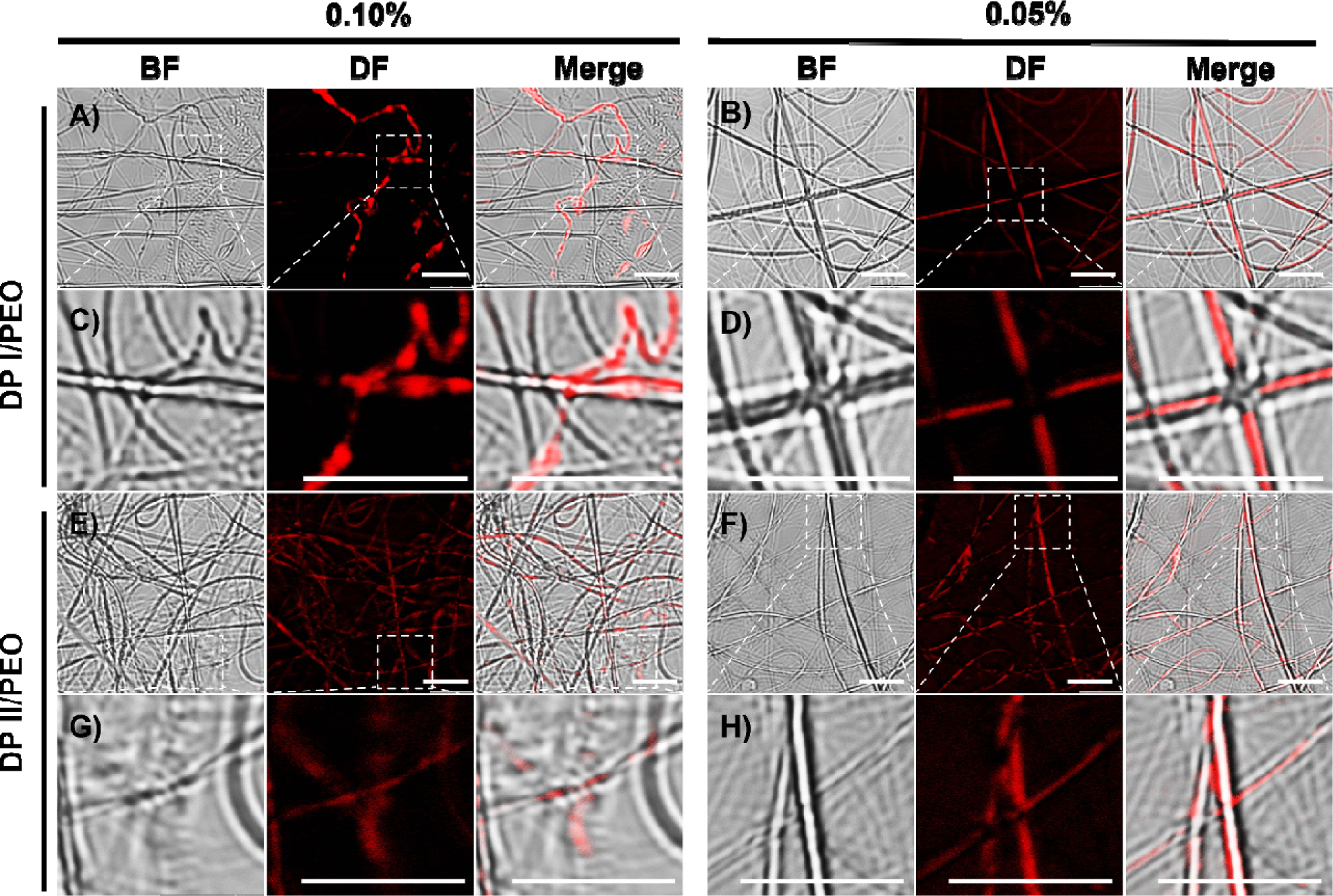
Morphology of PEO matrix electrospun with DPs: bright field (BF), dark field (DF), and merged images under fluorescence confocal microscopy. A-B) Blends of DP I with PEO matrix with 0.1% labeled (A) or 0.05% labeled peptide (B). C-D) Zoomed in areas in panels (A) and (B), respectively. E-F) Blends of DP II with PEO matrix with 0.1% labeled (A) or 0.05% labeled peptide (B). G-H) Zoomed in areas in panels (E) and (F), respectively. All scale bars are 5 μm.

### Cytotoxicity Evaluation and Antioxidant Activity

Historically, amyloids are aggregates of proteins considered pathogenic and toxic to neurons, but many amyloid materials in nature have been shown to have beneficial features.^40^ To confirm the potential of these materials for biomedical applications, both DPs were evaluated for toxicity to human cells using a lactate dehydrogenase (LDH) assay, and results are shown in Figure 7A. Following 24 hours of incubation with the cells, both DPs show no induced toxicity to human embryonic kidney cells (HEK 293) compared to control cells at all concentrations tested and no increase in toxicity compared to PEO. Thus, the presence of cationic segments did not impact the toxicity of the peptides, allowing applications, including wound dressings, to be considered for these highly biocompatible materials.

**Figure 7.**
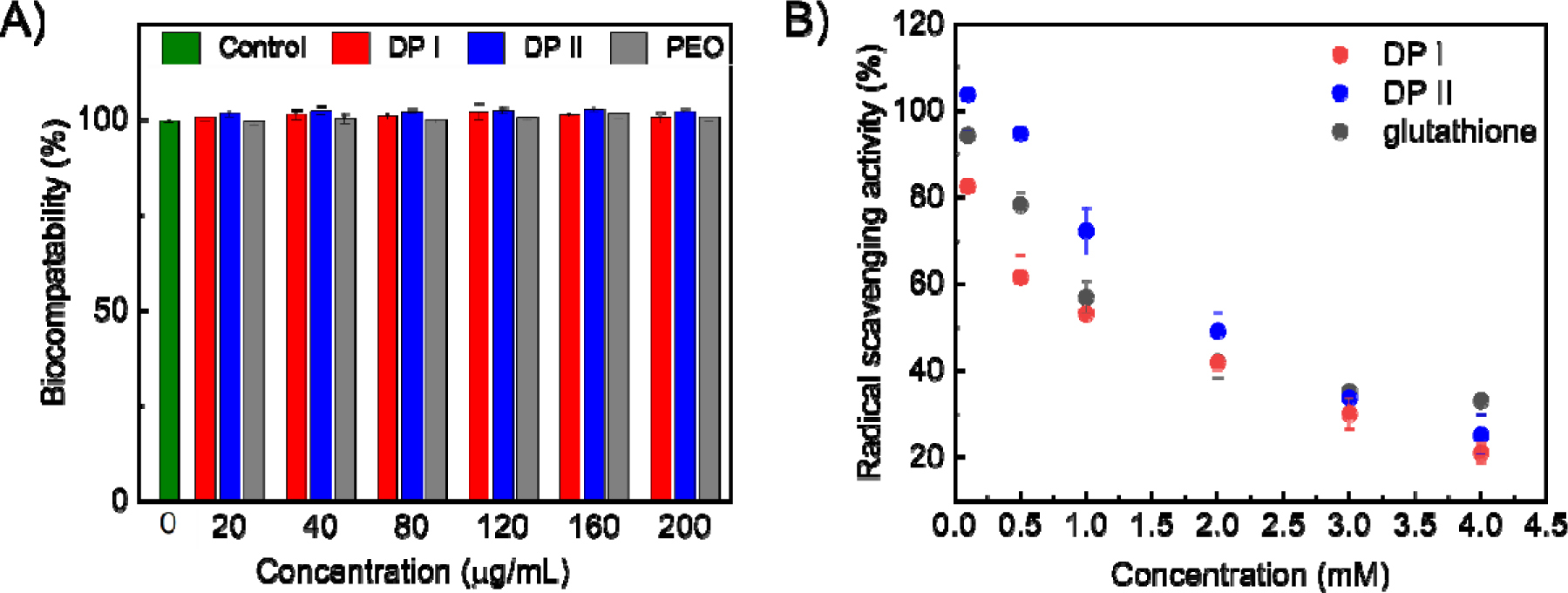
A) Biocompatibility evaluation of DPs. B) Antioxidation evaluation of DPs. Both DPs show no induced toxicity to human embryonic kidney cells and good antioxidant activity.

Free radicals accumulating at wound sites cause oxidative stress, which leads to lipid peroxidation, DNA damage, and enzyme inactivation, thereby impairing wound healing.^41^ The antioxidant activity of DPs was evaluated by comparing them to glutathione (the positive control) to explore their potential benefits as wound dressing materials. At low concentrations (0.1, 0.5, and 1 mM), DP I showed higher antioxidative activity than DP II at a statistically significant level, but no significant difference was observed at higher concentrations. DP I contains lysine and tryptophan, while DP II includes lysine, tryptophan, arginine, and methionine. Despite having more reductive pendant groups and theoretically stronger radical scavenging potential, DP II’s globular structures likely trap these pendant groups in the core, reducing their exposure on the surface and limiting their effectiveness. This comparison indicates that the higher-order assembled structure can significantly impact the peptide antioxidative performance beyond what might be predicted from their sequence alone.^42^

## CONCLUSIONS

In this research, two decapeptides with fibrillar (DP I) and globular (DP II) morphologies were blended with PEO ethanol/water solutions for electrospinning, and their morphological, mechanical, rheological, and thermal properties were systematically evaluated. The incorporation of DPs notably enhanced the mechanical properties of the composite mats, and such physical reinforcement seems to result from a uniform distribution of the DPs rather than changes in the crystallinity of the PEO matrix or nanofiber bulk alignment. The improved mechanical properties are impressive considering the thinner nanofibers in the composite mats. The lowered diameter in the amyloid containing fibers is attributed to increased conductivity of the electrospinning solution imparted by the zwitterionic nature of the peptides. Post-processing analysis revealed that both decapeptides retained their secondary structures within the DP/PEO blends, indicating stability of DPs during electrospinning. Despite these similarities, DPs show subtle but distinct differences when blended with PEOs, which seem to correlate with their secondary structure and fibril-forming propensities. Low cytotoxicity and strong antioxidant activity of DPs underscore their potential for safe and effective use in the biomedical field. These findings highlight the intricate connections between peptide structural features, DP/PEO blend properties, and nanocomposite mat processing. This study also demonstrates the strong potential of using peptide-polymer blends for developing advanced biomaterials with applications in biotechnology, particularly in areas requiring enhanced mechanical properties and bioactivity.

## EXPERIMENTAL SECTION

### Materials

Wang polystyrene resin, Fmoc protected amino acids, and ethyl cyanoglyoxlate-2-oxime (Oxyma) were purchased from CEM peptides. Polyethylene oxide (average *M*_v_∼1,000,000), 1,2-ethanedithiol (EDT), deuterated ethanol (C_2_D_5_OD) and deuterium oxide (D_2_O) were purchased from Sigma Aldrich Corporation ^43^. Dichloromethane (DCM), diethyl ether, trifluoroacetic acid (TFA), dimethylformamide (DMF), ethanol, acetonitrile (MeCN), water (H_2_O, HPLC grade), Dulbecco’s modified eagle medium (DMEM), heat inactivated fetal bovine serum (FBS), penicillin-streptomycin, and CyQUANT lactate dehydrogenase (LDH) assay were all purchased from ThermoFisher ^43^. Diisopropylcarbodiimide (DIC) and triisopropylsilane ^37^ were purchased from Acros Chemicals ^43^. 1,1-Diphenyl-2-picrylhydrazyl (DPPH) was purchased from Cayman Chemical Company. Sterile tissue treated 96-well plates were obtained from CellTreat ^43^. All chemicals were used without further purification.

### Peptide Synthesis and Purity Determination

Peptides were synthesized on a Liberty Blue 2.0 automated peptide synthesizer (CEM) through standard 9-fluorenyl methoxycarbonyl (Fmoc)-based solid phase peptide synthesis. Peptide synthesis was performed at 0.25 mmol scale using Fmoc-Asp(OtBu)-Wang Resin (LL) and Fmoc-Lys-Wang Resin (LL), for DP I and DP II, respectively. (0.3mmol/g loading, 100-200 mesh). Deprotection of Fmoc protecting groups was carried out using 20 v/v% piperidine in DMF. Each amino acid addition was carried out using Fmoc-protected amino acids (0.2 M), DIC (1M), and Oxyma (1 M) in DMF. After the final Fmoc deprotection, the resin beads were washed 3x using DCM. The peptide then underwent global deprotection and cleavage from the resin beads through gentle shaking in TFA/TIS/H_2_O/EDT (95: 2.5:2.5:2.5) cleavage cocktail for 3 hours at room temperature. Peptides were then precipitated in cold diethyl ether and collected via centrifugation. The peptide pellet was then resuspended in diethyl ether and chilled for four hours. It was recentrifuged, and the diethyl ether supernatant was decanted from the peptide pellet, which was allowed to air dry. Crude peptides were purified on a Prodigy preparative reverse-phase HPLC (CEM) with a H_2_O/MeCN gradient (containing 0.1% TFA). The mass and identity of the eluting fractions containing the desired DP peptides were confirmed using electrospray ionization ^27^-mass spectrometry (MS) on a Thermo Scientific Orbitrap Exploris™ 240. The purity was determined using liquid chromatography-mass spectrometry (LC-MS), shown in Figures S1 and S2. Labeling of decapeptides was carried out by incubating decapeptides (DP I and DP II) with 0.5x of fluorescent dye, HiLyte™ Fluor 647 succinimidyl ester *(AnaSpec Inc)* for 12h at 4 °C. For the reactions, 0.1% and 0.05% fluorescent dye were used with the unlabeled decapeptides, respectively.

### Solution Fourier Transform Infrared (FTIR) Spectroscopy

DP samples were prepared at 2.5 mg/mL by dissolving the dry peptides in C_2_D_5_OD: D_2_O mixture (v/v = 3/1) and incubated at room temperature for 48 hours. FTIR data was obtained at room temperature using an Agilent Cary 630 FTIR instrument. Samples were scanned for 1024 runs at 8 cm^−1^ resolution. Peak deconvolution was done with OriginLab 8.0 using the Gaussian algorithm.

### Atomic Force Microscopy (AFM)

AFM samples were prepared following a previously published procedure.^44^ Freshly cleaved mica substrates were first treated with 150 μL of APTES solution (500 μL of 3-aminopropyltriethoxysilane in 50 mL of 1 mM acetic acid) for 20 min. The APTES solution was decanted, and the surfaces were rinsed thrice with 150 μL H_2_O. The substrates were dried under a stream of N_2_ and stored in the desiccator for one h. Peptide solution (150 μL) was deposited onto the amine-treated mica substrates for 30 min to adsorb the peptide. The protein solution was then decanted, and the samples were rinsed three times with 150 μL H_2_O. The samples were dried under a stream of N_2_ and stored in the desiccator until imaging. All AFM experiments were performed using a Dimension Icon atomic force microscope (Bruker). AFM scanning was conducted using NanoScope 8.15r3sr9 software, and the images were analyzed using NanoScope Analysis 1.50 software. Imaging was performed using a sharp silicon nitride cantilever (RTESPA-300, nominal tip radius 8 nm; nominal resonance frequency of 300 kHz; nominal spring constant of 40 N/m) and a standard probe holder under ambient conditions with 512 × 512 data point resolution.

### PEO/Peptide Blend preparation

Table 4 summarizes the composition of each blend. The decapeptide was dissolved in ethanol/H_2_O mixture (v/v = 3/1) and incubated at room temperature for 48 hours before mixing with PEO. The final electrospinning solution was kept at a total PEO concentration of 3% (w/v) in 3:1 ethanol/water with different peptide concentrations (1.25, 2.50, and 5.00 mg/mL).

**Table 4.** Composition of PEO/DP blends.

| sample ID | DP | DP conc. (mg/mL) | PEO conc. (wt %) | PEO: DP |
| --- | --- | --- | --- | --- |
| PEO-3 | - | - | 3 | 100.0:0.0 |
| 3-I-1.25 | DP I | 1.25 | 3 | 96.0:4.0 |
| 3-I-2.5 | DP I | 2.50 | 3 | 92.3:7.7 |
| 3-I-5 | DP I | 5.00 | 3 | 85.5:14.2 |
| 3-II-1.25 | DP II | 1.25 | 3 | 96.0:4.0 |
| 3-II-2.5 | DP II | 2.50 | 3 | 92.3:7.7 |
| 3-II-5 | DP II | 5.00 | 3 | 85.5:14.2 |

### Solution Viscosity

Viscosity measurements were performed using a strain-controlled ARES rheometer with a cone-and-plate geometry (25 mm and 0.997 rad, or 50 mm and 0.499 rad). A steady rate sweep test was conducted at room temperature with a rate from 0.1 to 1000 s^−1^.

### Fabrication of PEO/Decapeptide Mats

Electrospun mats were fabricated using a Lab Mate 30kV Electrospinning Machine (Spruce Science, CA, USA). Fibers were collected on a home-built aluminum drum collector with a collection width of 0.8 cm and a drum diameter of 7.5 cm, with a needle-collector distance of 22.5 cm and a rotation speed of 60 rpm, shown in Figure S3. The screening fabrication conditions were a 21G needle, 20 kV, and a flow rate of 0.25 mL/h.

### Attenuated Total Reflectance Fourier Transform Infrared (ATR-FTIR) Spectroscopy

FTIR spectra were obtained using a Thermo Scientific Nicolet iS50 FT-IR in ATR mode. Reported spectra are an average of 128 scans at a resolution of 1 cm^−1^. FTIR data was normalized and analyzed using Origin Lab 8.0.

### Scanning Electron Microscopy (SEM)

Analysis of the mat morphology was done using a Zeiss Sigma VP field-emission SEM with Thermo System 7 Energy Dispersive X-ray Spectroscopy (EDS) and Wavelength Dispersive X-ray Spectrometry (WDS) X-ray detectors (Thermo-Fisher Scientific, Waltham, MA). Electrospinning samples were carbon-coated using a Cressington Carbon Coater (coating pressure < 0.01 mbar, coating duration ten seconds repeated ten times for each sample). The fiber morphology was analyzed using ImageJ Analysis (ImageJ, National Institute of Health, MD).

### Differential Scanning Calorimetry

The crystalline changes of electrospun fibers were monitored using a TA Instruments Discovery DSC 250 using aluminum hermetic pans. DSC thermograms were obtained using heat/cool/heat cycles from −80 to 100 °C with a heating ramp rate of 10 °C min^−1^ in all cycles. The degree of crystallinity (*X*) was determined using Equation 1:

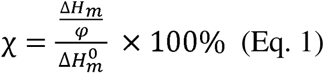

where 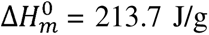 is the melting enthalpy when crystallinity of PEO is 100%, Δ*H_m_* the melting enthalpy derived from the DSC curves, and φ is the weight fraction of PEO in the composite mat, (control: φ = 1; 1.25 mg/mL loading, φ = 0.96; 2.5 mg/mL loading, φ = 0.923; 5 mg/mL loading, φ = 0.857).^45^

### Solution Conductivity

The conductivity of the DP ethanol/water solution was measured using an Accumet XL 500 Dual Channel pH/mV/Ion/Conductivity meter. The measurement was conducted in single-point mode, with a cell constant of 1.000/cm. Calibration was performed using a conductivity standard of 147 μmhos/cm.

### Tensile test

Tensile testing was performed using a Mark-10 EasyMESUR Test (Model F105) with a 25 N load cell. Electrospun mat samples were peeled from the collector, cut into strips with a length of 3.5 cm and width of 0.8 cm, and affixed to a paper frame using scotch tape, shown in Figure S4A. Samples were tested in at a pull-off rate of 13 mm/min. Mat thickness was estimated based on the mass normalization method.^46–48^ The stress was calculated using Equation 2:

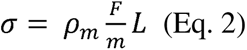

where *p_m_* is the material bulk density, *m* is the specimen mass, *L* is the specimen initial length, *F* is the force, and σ is the stress. For PEO and composite mats, *p_m_* is 1.13 g/cm^3^, as reported in the technical datasheet, and we assumed that low peptide loading would not significantly change the composite mat bulk density. Tests were run in triplicate and samples were collected from two independent mat fabrications, n = 6.

### Fluorescence Microscopy and Sample Preparation

Fluorescence microscopic images of decapeptide fibers were obtained using Leica SP8 confocal microscope at 100X magnification. All images were processed using Affinity Designer. The labeled peptide (0.1%) was incubated with unlabeled peptide in a 3:1 ethanol/water mixture for 48 h prior to blending with PEO stock solution. The collector was changed to a flat aluminum plate with a glass cover (#1 micro cover glass VWR Scientific) placed on the top. After one hour of random fiber collection in the dark, the glass coverslip was carefully placed onto a plain microscope slide (Fisherbrand) to sandwich the fibers between the slide and coverslip. All glass substrates were cleaned with ethanol and wiped with lens paper prior to collection. The sample assembly was secured using instant dry topcoat (Sally Hansen) and stored in the dark prior to fluorescence imaging, shown in Figure S5.

### Cytotoxicity and Antioxidation Evaluation

Human embryonic kidney cells (HEK293) were grown to 90% confluence in tissue culture-treated polystyrene flasks in an incubator held at 37 °C with 5% CO_2_. Cells were grown in DMEM supplemented with 10% FBS and 1% penicillin-streptomycin. HEK293 cells were seeded in a 96-well plate (1 × 105 cells/mL, 100 µL volume per well) and were incubated at 37 °C and 5% CO_2_ overnight to allow cells to adhere. DP I, DP II, and PEO were dissolved in supplemented DMEM at 1 mg/mL to create a stock solution, followed by serial dilutions with DMEM to obtain the final treatment concentration with cells. Each concentration was assessed in triplicate. Nuclease-free water was used as a spontaneous LDH control, and Triton X-100 was used as a positive control for 100 % cytotoxicity (i.e., complete LDH release). Plates were incubated for 24 h at 37 °C and 5% CO_2_ before collecting media to assess LDH release with the CyQUANT LDH assay following the manufacturers protocols.

DPPH solution (100 uM, in ethanol/water (V/V = 3:1)) was freshly prepared before use. Glutathione, a known antioxidant, was selected as the positive control. The DPPH radical-scavenging activity was determined using the method of Santos et al.^43^ DPPH solution (50 uL) was mixed with 50 uL analyte solutions (100, 500,1000, 2000, 3000, 4000 uM). The mixture was kept in a dark environment at room temperature for 30 min. The normalized DPPH intensity was calculated using Eq 3:

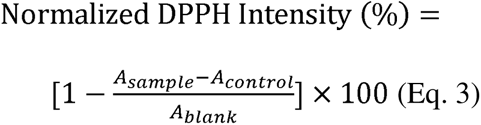

where A_sample_ is the absorbance value of the 50 uL sample mixed with 50 uL DPPH solution, A_control_ is the absorbance value of 50 uL sample solution mixed with 50 uL ethanol/water solution, and A_blank_ is the absorbance value of 50 uL DPPH solution mixed with 50 uL ethanol/water solution alone.^49^ Lower normalized intensity means more DPPH radicals are quenched, indicating stronger antioxidant activity. Statistical significance of differences in antioxidant activity at each concentration was determined by one-way ANOVA, p < 0.05.

## ASSOCIATED CONTENT

Figure S 1 A) chemical structure and molecular weight of DPI. B) LC-MS trace of crude DPI and C) corresponding crude ESI-mass spectra. D) LC-MS trace of pure DPI and E) corresponding pure ESI-mass spectra.

Figure S 1 A) chemical structure and molecular weight of DPII. B) LC-MS trace of crude DPII and C) corresponding crude ESI-mass spectra. D) LC-MS trace of pure DPII and E) corresponding pure ESI-mass spectra.

Figure S 3 A) Electrospinning setup. B) The collecting station can move horizontally while the collector rotates. C) A picture of the home-built collector.

Figure S 2 A) The paper frame used to prepare tensile bars. B) A snapshot of the tensile test. Figure S 3 Aluminum/glass cover as collector (left) and sandwiched sample for imaging.

Table S1 Conductivity of DP solutions.

## AUTHOR INFORMATION

### Author Contributions

The manuscript was written through contributions of all authors. All authors have given approval to the final version of the manuscript.

## ACKNOWLEDGEMENTS

We gratefully acknowledge financial support for this work from the National Science Foundation (NSF) (BMAT 2208349). Additionally, TDC acknowledges funding support from the National Institutes of Health (NIH) and the National Institute of Biomedical Imaging and Bioengineering (NIBIB R03EB033704). The cell work and fluorescence microscopy were supported by the Mississippi INBRE, funded by an Institutional Development Award (IDeA) from P20GM103476 from the NIH National Institutes of General Medical Sciences. Additionally, the authors acknowledge support from the National Science Foundation (Award number: 2319932) for mass spectrometry instrumentation utilized in this study. LCMS was performed at the Peptide Synthesis Core Facility of the Simpson Querrey Institute for BioNanotechnology at Northwestern University. This facility has current support from the Soft and Hybrid Nanotechnology Experimental (SHyNE) Resource (NSF ECCS-2025633).

## REFERENCES

(1) Rambaran, R. N.; Serpell, L. C. Amyloid fibrils. Prion 2008, 2 (3), 112–117. DOI: 10.4161/pri.2.3.7488.

(2) Balistreri, A.; Goetzler, E.; Chapman, M. Functional amyloids are the rule rather than the exception in cellular biology. Microorganisms 2020, 8 (12). DOI: 10.3390/microorganisms8121951.

(3) Otzen, D.; Riek, R. Functional amyloids. . Cold Spring Harb. Perspect. Biol. 2019, (1943-0264 (Electronic)).

(4) Schleeger, M.; vandenAkker, C. C.; Deckert-Gaudig, T.; Deckert, V.; Velikov, K. P.; Koenderink, G.; Bonn, M. Amyloids: From molecular structure to mechanical properties. Polymer 2013, 54 (10), 2473–2488. DOI: 10.1016/j.polymer.2013.02.029.

(5) Prosswimmer, T.; Heng, A.; Daggett, V. Mechanistic insights into the role of amyloid-β in innate immunity. Sci. Rep. 2024, 14 (1), 5376. DOI: 10.1038/s41598-024-55423-9.

(6) Sønderby, T. V.; Najarzadeh, Z.; Otzen, D. E. Functional bacterial amyloids: Understanding fibrillation, regulating biofilm fibril formation and organizing surface assemblies. In Molecules, 2022; Vol. 27.

(7) Rasmussen, H. Ø.; Kumar, A.; Shin, B.; Stylianou, F.; Sewell, L.; Xu, Y.; Otzen, D. E.; Pedersen, J. S.; Matthews, S. J. Fapa is an intrinsically disordered chaperone for pseudomonas functional amyloid fapc. J. Mol. Biol. 2023, 435 (2), 167878. DOI: 10.1016/j.jmb.2022.167878.

(8) Peña-Díaz, S.; Olsen, W. P.; Wang, H.; Otzen, D. E. Functional amyloids: The biomaterials of tomorrow? Adv. Mater. 2024, 36 (18), 2312823. DOI: 10.1002/adma.202312823.

(9) Van Houdt, R.; Michiels, C. W. Role of bacterial cell surface structures in escherichia coli biofilm formation. Res. Microbiol. 2005, 156 (5), 626–633. DOI: 10.1016/j.resmic.2005.02.005.

(10) Kikuchi, T.; Mizunoe, Y.; Takade, A.; Naito, S.; Yoshida, S.-i. Curli fibers are required for development of biofilm architecture in escherichia coli k-12 and enhance bacterial adherence to human uroepithelial cells. Microbiol. Immunol. 2005, 49 (9), 875–884. DOI: 10.1111/j.1348-0421.2005.tb03678.x.

(11) Wang, A.; Keten, S. Adhesive behavior and detachment mechanisms of bacterial amyloid nanofibers. npj Comput. Mater. 2019, 5 (1), 29. DOI: 10.1038/s41524-019-0154-7.

(12) Shanmugam, N.; Baker, M. O. D. G.; Ball, S. R.; Steain, M.; Pham, C. L. L.; Sunde, M. Microbial functional amyloids serve diverse purposes for structure, adhesion and defence. Biophys. Rev. 2019, 11 (3), 287–302. DOI: 10.1007/s12551-019-00526-1.

(13) Chen, D.; Liu, X.; Chen, Y.; Lin, H. Amyloid peptides with antimicrobial and/or microbial agglutination activity. Appl. Microbiol. Biotechnol. 2022, 106 (23), 7711–7720. DOI: 10.1007/s00253-022-12246-w.

(14) Riek, R. The three-dimensional structures of amyloids. . Cold Spring Harb. Perspect. Biol. 2017, (1943-0264 (Electronic)).

(15) Knowles, T. P.; Fitzpatrick, A. W.; Meehan, S.; Mott, H. R.; Vendruscolo, M.; Dobson, C. M.; Welland, M. E. Role of intermolecular forces in defining material properties of protein nanofibrils. Science 2007, 318 (5858), 1900–1903. DOI: 10.1126/science.1150057.

(16) Wang, W.; Azizyan, R. A.; Garro, A.; Kajava, A. V.; Ventura, S. Multifunctional amyloid oligomeric nanoparticles for specific cell targeting and drug delivery. Biomacromolecules 2020, 21 (10), 4302–4312. DOI: 10.1021/acs.biomac.0c01103.

(17) Diaz, C.; Missirlis, D. Amyloid-based albumin hydrogels. Adv. Healthcare Mater. 2023, 12 (7), 2201748. DOI: 10.1002/adhm.202201748.

(18) Das, S.; Jacob, R. S.; Patel, K.; Singh, N.; Maji, S. K. Amyloid fibrils: Versatile biomaterials for cell adhesion and tissue engineering applications. Biomacromolecules 2018, 19 (6), 1826–1839. DOI: 10.1021/acs.biomac.8b00279.

(19) Peydayesh, M.; Suter, M. K.; Bolisetty, S.; Boulos, S.; Handschin, S.; Nyström, L.; Mezzenga, R. Amyloid fibrils aerogel for sustainable removal of organic contaminants from water. Adv. Mater. 2020, 32 (12), 1907932. DOI: 10.1002/adma.201907932.

(20) Bolisetty, S.; Reinhold, N.; Zeder, C.; Orozco, M. N.; Mezzenga, R. Efficient purification of arsenic-contaminated water using amyloid–carbon hybrid membranes. Chem. Commun. 2017, 53 (42), 5714–5717. DOI: 10.1039/C7CC00406K.

(21) Jia, X.; Peydayesh, M.; Huang, Q.; Mezzenga, R. Amyloid fibril templated mof aerogels for water purification. Small 2022, 18 (4), 2105502. DOI: 10.1002/smll.202105502.

(22) Men, D.; Guo, Y.-C.; Zhang, Z.-P.; Wei, H.-p.; Zhou, Y.-F.; Cui, Z.-Q.; Liang, X.-S.; Li, K.; Leng, Y.; You, X.-Y.;, et al. Seeding-induced self-assembling protein nanowires dramatically increase the sensitivity of immunoassays. Nano Lett. 2009, 9 (6), 2246–2250. DOI: 10.1021/nl9003464.

(23) Saldanha, D. J.; Abdali, Z.; Modafferi, D.; Janfeshan, B.; Dorval Courchesne, N.-M. Fabrication of fluorescent ph-responsive protein–textile composites. Sci. Rep. 2020, 10 (1), 13052. DOI: 10.1038/s41598-020-70079-x.

(24) Li, T.; Kambanis, J.; Sorenson, T. L.; Sunde, M.; Shen, Y. From fundamental amyloid protein self-assembly to development of bioplastics. Biomacromolecules 2024, 25 (1), 5–23. DOI: 10.1021/acs.biomac.3c01129.

(25) Peydayesh, M.; Bagnani, M.; Mezzenga, R. Sustainable bioplastics from amyloid fibril-biodegradable polymer blends. ACS Sustainable Chem. Eng. 2021, 9 (35), 11916–11926. DOI: 10.1021/acssuschemeng.1c03937.

(26) Bagnani, M.; Ehrengruber, S.; Soon, W. L.; Peydayesh, M.; Miserez, A.; Mezzenga, R. Rapeseed cake valorization into bioplastics based on protein amyloid fibrils. Adv. Mater. Technol. 2023, 8 (3), 2200932. DOI: 10.1002/admt.202200932.

(27) Rubin, D. J.; Nia, H. T.; Desire, T.; Nguyen, P. Q.; Gevelber, M.; Ortiz, C.; Joshi, N. S. Mechanical reinforcement of polymeric fibers through peptide nanotube incorporation. Biomacromolecules 2013, 14 (10), 3370–3375. DOI: 10.1021/bm4008293.

(28) Kaur, G.; Kumari, S.; Saha, P.; Ali, R.; Patil, S.; Ganesh, S.; Verma, S. Selective cell adhesion on peptide–polymer electrospun fiber mats. ACS Omega 2019, 4 (2), 4376–4383. DOI: 10.1021/acsomega.8b03494.

(29) Abernathy, H. G.; Saha, J.; Kemp, L. K.; Wadhwani, P.; Clemons, T. D.; Morgan, S. E.; Rangachari, V. De novo amyloid peptides with subtle sequence variations differ in their self-assembly and nanomechanical properties. Soft Matter 2023, 19 (27), 5150–5159. DOI: 10.1039/D3SM00604B.

(30) Topuz, F.; Abdulhamid, M. A.; Holtzl, T.; Szekely, G. Nanofiber engineering of microporous polyimides through electrospinning: Influence of electrospinning parameters and salt addition. Mater. Des. 2021, 198, 109280. DOI: 10.1016/j.matdes.2020.109280.

(31) Qin, X.-H.; Yang, E.-L.; Li, N.; Wang, S.-Y. Effect of different salts on electrospinning of polyacrylonitrile (pan) polymer solution. J. Appl. Polym. Sci. 2007, 103 (6), 3865–3870. DOI: 10.1002/app.25498.

(32) Zong, X.; Kim, K.; Fang, D.; Ran, S.; Hsiao, B. S.; Chu, B. Structure and process relationship of electrospun bioabsorbable nanofiber membranes. Polymer 2002, 43 (16), 4403–4412. DOI: 10.1016/S0032-3861(02)00275-6.

(33) Zhou, C.; Chu, R.; Wu, R.; Wu, Q. Electrospun polyethylene oxide/cellulose nanocrystal composite nanofibrous mats with homogeneous and heterogeneous microstructures. Biomacromolecules 2011, 12 (7), 2617–2625. DOI: 10.1021/bm200401p.

(34) Pittarate, C.; Yoovidhya, T.; Srichumpuang, W.; Intasanta, N.; Wongsasulak, S. Effects of poly(ethylene oxide) and zno nanoparticles on the morphology, tensile and thermal properties of cellulose acetate nanocomposite fibrous film. Polym. J. 2011, 43 (12), 978–986. DOI: 10.1038/pj.2011.97.

(35) Samal, S.; Kim, S.; Kim, H. Effects of filler size and distribution on viscous behavior of glass composites. J. Am. Ceram. Soc. 2012, 95 (5), 1595–1603. DOI: 10.1111/j.1551-2916.2012.05181.x.

(36) Kataoka, T.; Kitano, T.; Sasahara, M.; Nishijima, K. Viscosity of particle filled polymer melts. Rheol. Acta. 1978, 17 (2), 149–155. DOI: 10.1007/BF01517705.

(37) Mutiso, R. M.; Winey, K. I. Electrical conductivity of polymer nanocomposites. In Polymer science: A comprehensive reference, Matyjaszewski, K., Möller, M. Eds.; Elsevier, 2012; pp 327–344.

(38) Sim, L. H.; Gan, S. N.; Chan, C. H.; Yahya, R. Atr-ftir studies on ion interaction of lithium perchlorate in polyacrylate/poly(ethylene oxide) blends. Spectrochim. Acta A Mol. Biomol. Spectrosc. 2010, 76 (3), 287–292. DOI: 10.1016/j.saa.2009.09.031.

(39) Pielichowska, K.; Głowinkowski, S.; Lekki, J.; Biniaś, D.; Pielichowski, K.; Jenczyk, J. Peo/fatty acid blends for thermal energy storage materials. Structural/morphological features and hydrogen interactions. Eur. Polym. J. 2008, 44 (10), 3344–3360. DOI: 10.1016/j.eurpolymj.2008.07.047.

(40) Yakupova, E. I.; Bobyleva, L. G.; Shumeyko, S. A.; Vikhlyantsev, I. M.; Bobylev, A. G. Amyloids: The history of toxicity and functionality. Biology 2021, 10 (5). DOI: 10.3390/biology10050394.

(41) Zhao, Y.; Li, X.; Sun, N.; Mao, Y.; Ma, T.; Liu, X.; Cheng, T.; Shao, X.; Zhang, H.; Huang, X.;, et al. Injectable double crosslinked hydrogel-polypropylene composite mesh for repairing full-thickness abdominal wall defects. Adv. Healthcare Mater. 2024, 13 (15), 2304489. DOI: 10.1002/adhm.202304489.

(42) Zou, T.-B.; He, T.-P.; Li, H.-B.; Tang, H.-W.; Xia, E.-Q. The structure-activity relationship of the antioxidant peptides from natural proteins. In Molecules, 2016; Vol. 21.

(43) Santos, J. S.; Alvarenga Brizola, V. R.; Granato, D. High-throughput assay comparison and standardization for metal chelating capacity screening: A proposal and application. Food Chem. 2017, 214, 515–522. DOI: 10.1016/j.foodchem.2016.07.091.

(44) Saha, J.; Ford, B. J.; Wang, X.; Boyd, S.; Morgan, S. E.; Rangachari, V. Sugar distributions on gangliosides guide the formation and stability of amyloid-β oligomers. Biophys. Chem. 2023, 300, 107073. DOI: 10.1016/j.bpc.2023.107073.

(45) Money, B. K.; Swenson, J. Dynamics of poly(ethylene oxide) around its melting temperature. Macromolecules 2013, 46 (17), 6949–6954. DOI: 10.1021/ma4003598.

(46) Maccaferri, E.; Cocchi, D.; Mazzocchetti, L.; Benelli, T.; Brugo, T. M.; Giorgini, L.; Zucchelli, A. How nanofibers carry the load: Toward a universal and reliable approach for tensile testing of polymeric nanofibrous membranes. Macromol. Mater. Eng. 2021, 306 (7), 2100183. DOI: 10.1002/mame.202100183.

(47) Munawar, M. A.; Schubert, D. W. Highly oriented electrospun conductive nanofibers of biodegradable polymers-revealing the electrical percolation thresholds. ACS Appl. Polym. Mater. 2021, 3 (6), 2889–2901. DOI: 10.1021/acsapm.0c01332.

(48) Maccaferri, E.; Ortolani, J.; Mazzocchetti, L.; Benelli, T.; Brugo, T. M.; Zucchelli, A.; Giorgini, L. New application field of polyethylene oxide: Peo nanofibers as epoxy toughener for effective cfrp delamination resistance improvement. ACS Omega 2022, 7 (27), 23189–23200. DOI: 10.1021/acsomega.2c01189.

(49) Yang, L.; Xing, Y.; Chen, R.; Ni, H.; Li, H.-H. Isolation and identification of antioxidative peptides from crocodile meat hydrolysates using silica gel chromatography. Sci. Rep. 2022, 12 (1), 13223. DOI: 10.1038/s41598-022-16009-5.

