## Supplementary material for "Amyloid peptide – synthetic polymer blends with enhanced mechanical and biological properties": supp data

**This PDF file includes:**

Supplementary Figure 1 to 5

Supplementary Table 1

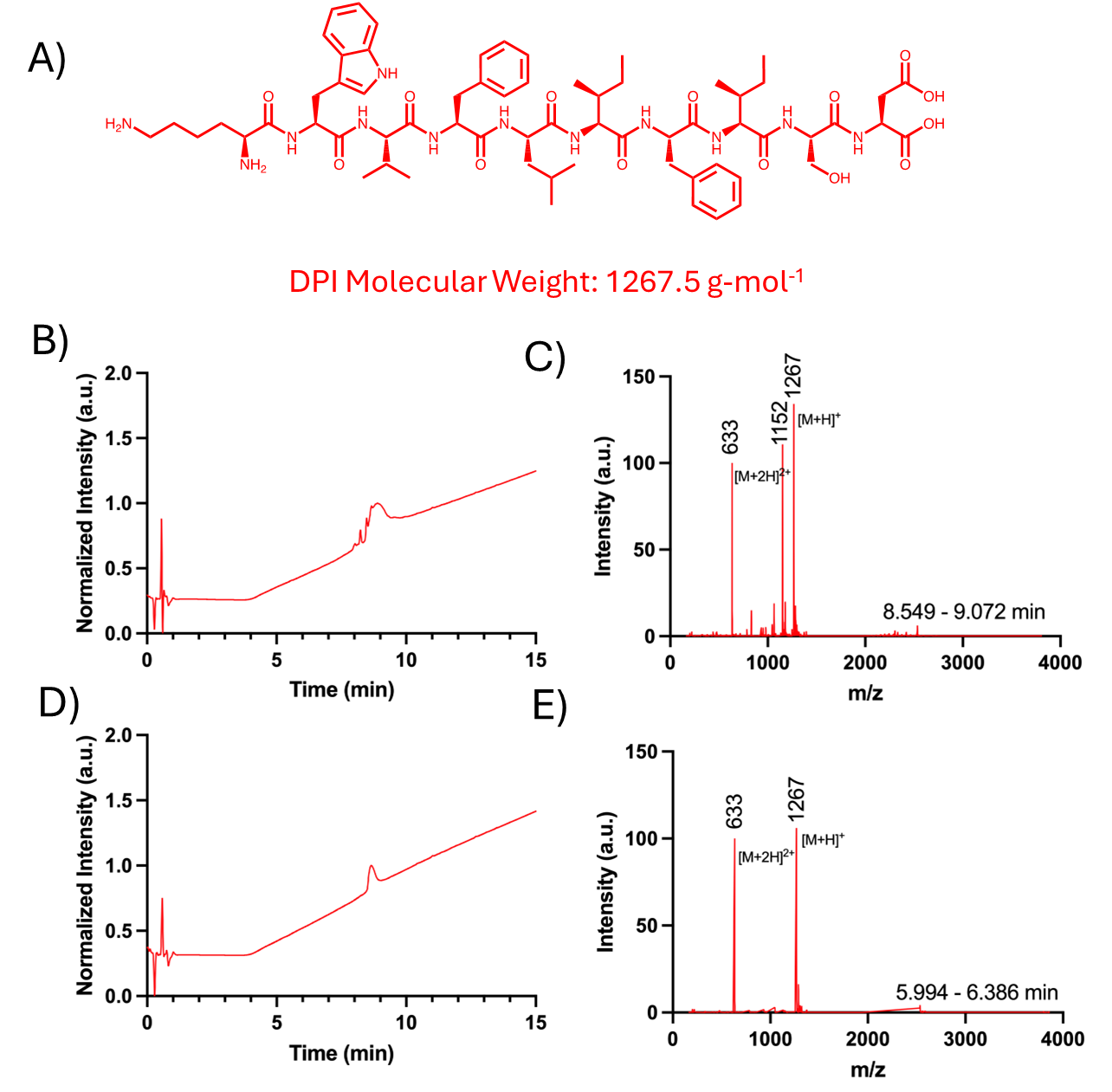

Figure S 1 A) chemical structure and molecular weight of DPI. B) LC-MS trace of crude DPI and C) corresponding crude ESI-mass spectra. D) LC-MS trace of pure DPI and E) corresponding pure ESI-mass spectra. LC-MS conditions: [peptide] = 1 mg/mL, loading solvent; H_2_O with 0.1% TFA (v/v), eluent; H_2_O-CH_3_CN gradient containing 0.1% HCOOH (v/v), column; Phenomenex Gemini 5 µm C18 110 Å LC column 150 x 1 mm. DP I is approximately 75% pure.

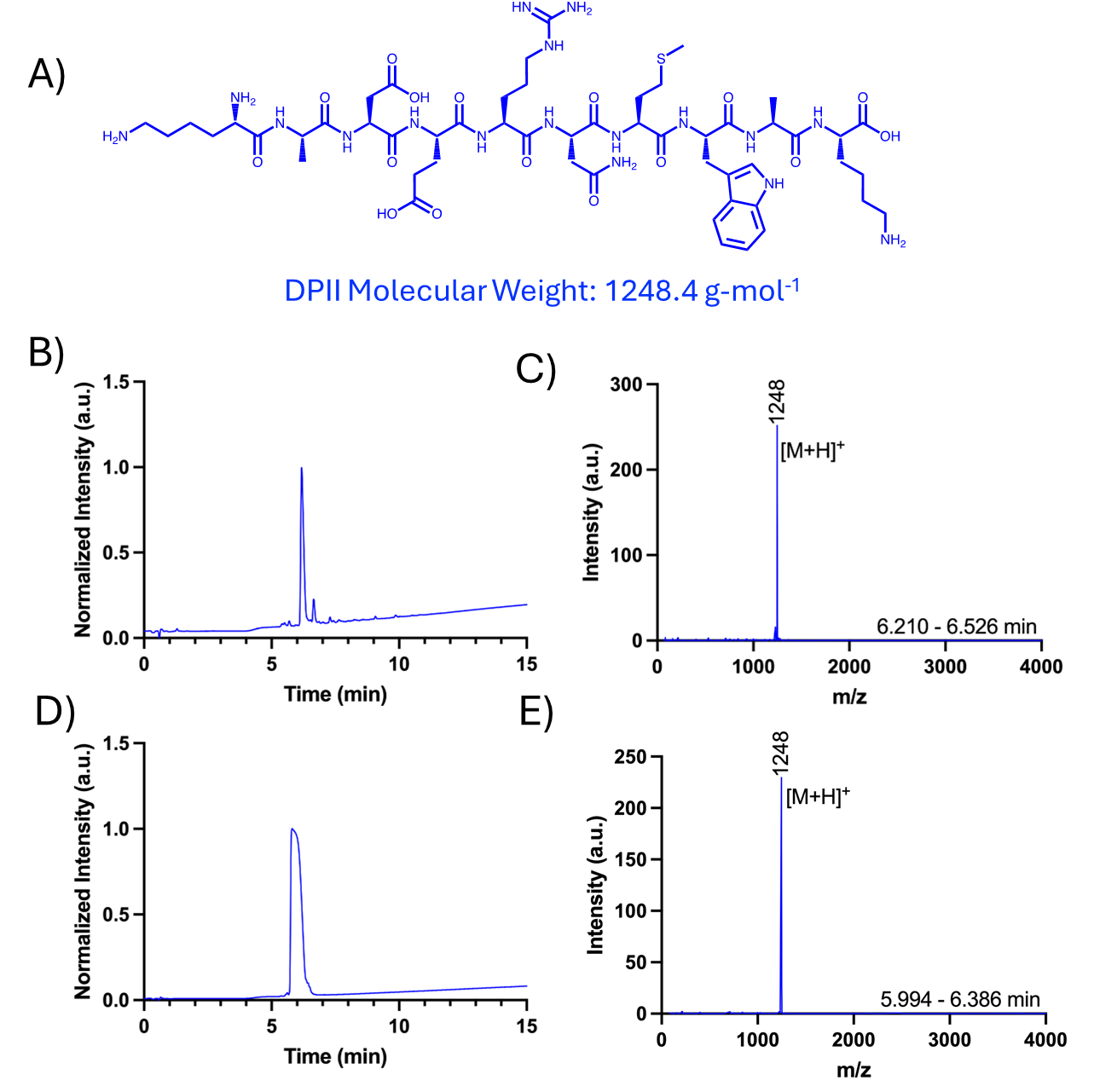

Figure S 2 A) chemical structure and molecular weight of DPII. B) LC-MS trace of crude DPII and C) corresponding crude ESI-mass spectra. D) LC-MS trace of pure DPII and E) corresponding pure ESI-mass spectra. LC-MS conditions: [peptide] = 1 mg/mL, loading solvent; H_2_O with 0.1% TFA (v/v), eluent; H_2_O-CH_3_CN gradient containing 0.1% HCOOH (v/v), column; Phenomenex Gemini 5 µm C18 110 Å LC column 150 x 1 mm. DP II is approximately 90% pure.

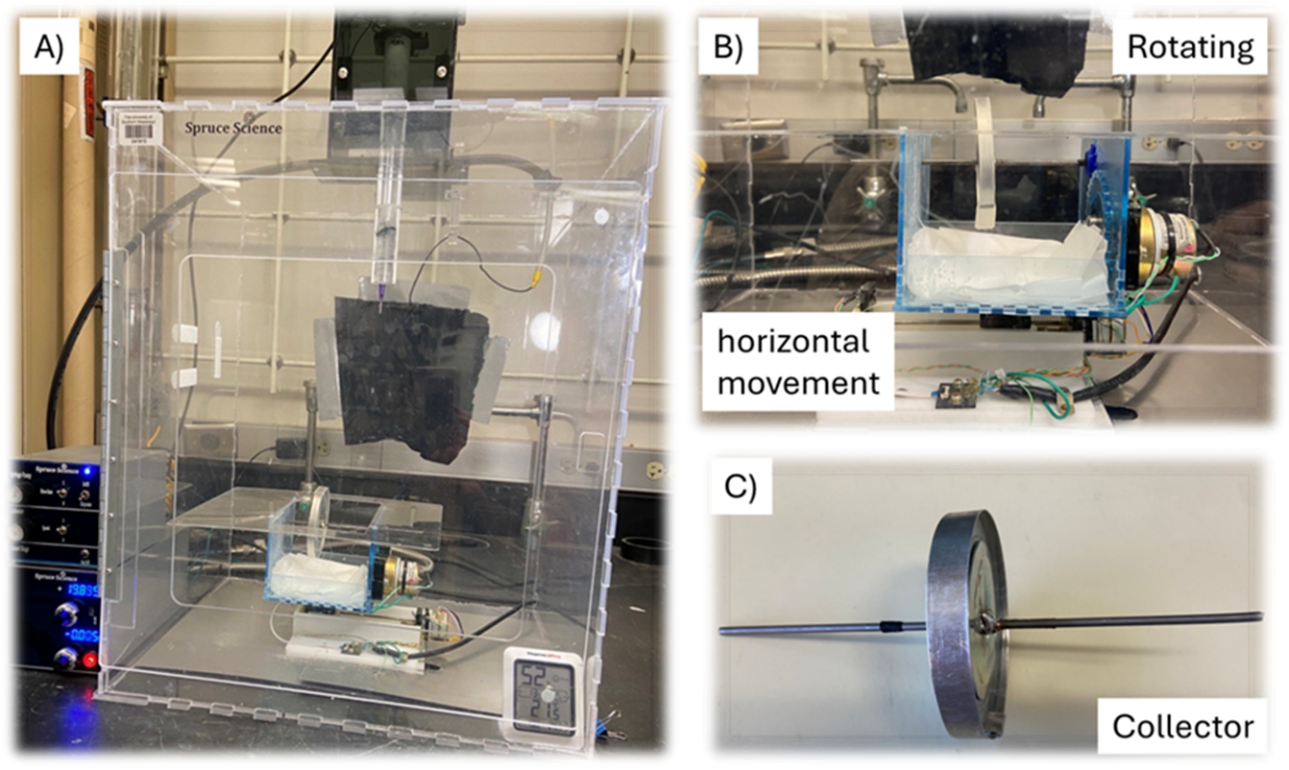

Figure S 3 A) Electrospinning setup. B) The collecting station can move horizontally while the collector rotates. C) A picture of the home-built collector. The collector will move horizontally and rotate simultaneously during collection.

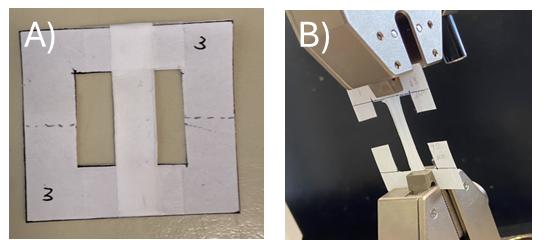

Figure S 4 A) The paper frame used to prepare tensile bars. B) A snapshot of the tensile test.

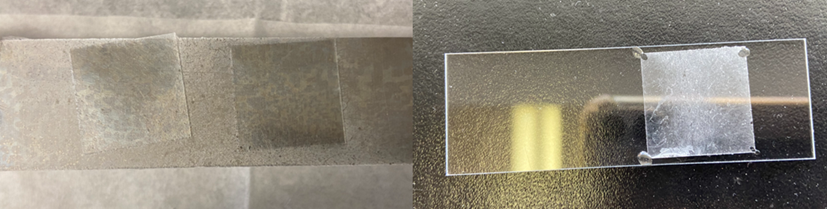

Figure S 5 Aluminum/glass cover as collector (left) and sandwiched sample for imaging (right).

Table S1 Conductivity of DP solutions.

|  | DP I solution (μS/cm) | DP II solution (μS/cm) |
| --- | --- | --- |
| 5.00 mg/mL | 269.8 ± 1.1 | 292.1 ± 2.4 |
| 2.50 mg/mL | 139.6 ± 0.5 | 184.6 ± 2.4 |
| 1.25 mg/mL | 78.8 ± 0.2 | 111.2 ± 1.1 |
